# Mechanisms of integrin αVβ5 clustering in flat clathrin lattices

**DOI:** 10.1101/427112

**Authors:** Alba Zuidema, Wei Wang, Maaike Kreft, Lisa te Molder, Liesbeth Hoekman, Onno B. Bleijerveld, Leila Nahidiaza, Hans Janssen, Arnoud Sonnenberg

## Abstract

This article highlights several molecular mechanisms that result in the assembly of integrin αVβ5-containing flat clathrin lattices in human keratinocytes.

**Abstract:** The family of integrin transmembrane receptors is essential for the normal function of multicellular organisms by facilitating cell-extracellular matrix adhesion. The vitronectin-binding integrin αVβ5 localizes to focal adhesions (FAs) as well as poorly characterized flat clathrin lattices (FCLs). Here we show that in human keratinocytes αVβ5 is predominant found in FCLs and that formation of the αVβ5-containing FCLs requires the presence of vitronectin as ligand, calcium, and the clathrin adaptor proteins ARH, Numb, and EPS15/EPS15L1. Integrin chimeras, containing the extracellular and transmembrane domains of β5 and the cytoplasmic domains of β1 or β3, almost exclusively localize in FAs. Interestingly, lowering actomyosin-mediated contractility promotes integrin redistribution to FLCs in an integrin tail-dependent manner, while increasing cellular tension favors αVβ5 clustering in FAs. Our findings strongly indicate that clustering of integrin αVβ5 in FCLs is dictated by the β5 subunit cytoplasmic domain, cellular tension, and recruitment of specific adaptor proteins to the β5 subunit cytoplasmic domains.

## Introduction

Cell adhesion to the surrounding extracellular matrix (ECM) is a basic requirement in multicellular organisms. Integrins are a major family of cell adhesion transmembrane receptors that are formed through heterodimerization of α and β subunits (Hynes, 1987). Integrins can assemble different types of cell-matrix adhesions, for instance by clustering in focal adhesions (FAs) and forming a mechanical link between the ECM and intracellular actin bundles (Geiger and Yamada, 2011; Jansen et al., 2017) or by connecting the ECM to the intracellular intermediate filament system in hemidesmosomes (de Pereda et al., 2009; Litjens et al., 2006). Other integrin-containing complexes include tetraspanin-enriched microdomains (Charrin et al., 2014; Hemler, 2005), fibrillar adhesions (FBs) (Geiger and Yamada, 2011), podosomes (Geiger and Yamada, 2011; Linder and Wiesner, 2016) and flat clathrin lattices (FCLs) (Grove et al., 2014; Lampe et al., 2016).

Depending on the combination of α and β subunits integrins bind different ECM components, including laminins, collagens, and fibronectin (Barczyk et al., 2010; Hynes, 2002). The RGD-binding integrins form a subset of integrins that recognize an arginine (R), glycine (G), aspartic acid (D) tri-peptide sequence present in ECM proteins such as fibronectin and vitronectin (Bodary and McLean, 1990; Charo et al., 1990; Cheresh et al., 1989; Cheresh and Spiro, 1987; Smith and Cheresh, 1990). Several RGD-binding integrins can bind the same ligands, but despite this, RGD-binding integrins can have distinct subcellular localization patterns. For example, in many cell types the integrin αVβ3 localizes exclusively in FAs, while α5β1 is present in both FAs and FBs. Additionally, in some cell types, these integrins are concentrated in podosomes. The integrin αVβ5 can be found in both FAs and clathrin-coated membrane domains (De Deyne et al., 1998; Leyton-Puig et al., 2017; Wayner et al., 1991).

The clathrin structures in which integrin αVβ5 clusters reside, have in later years been described as FCLs or clathrin “plaques” (Akisaka et al., 2008; Grove et al., 2014; Lampe et al., 2016; Leyton-Puig et al., 2017; Maupin and Pollard, 1983; Saffarian et al., 2009). These structures comprise large assemblies of clathrin that are distinct from the more dynamic clathrin-coated pits that play an active role in the selective internalization of membrane-bound proteins through a process known as clathrin-mediated endocytosis (CME) (Kaksonen and Roux, 2018; Lampe et al., 2016). The physiological relevance of FCLs is not completely understood. They could be involved in CME by providing stable platforms for the recruitment of cargo (Grove et al., 2014) and/or play a role in cell adhesion (Batchelder and Yarar, 2010; Elkhatib et al., 2017; Saffarian et al., 2009).

Since RGD-binding integrins are mainly found to reside in FAs and associate with the actin cytoskeleton, we wondered which processes drive the clustering of integrins in other cell adhesion structures. In the present study, we unravel different mechanisms that lead to integrin αVβ5 clustering in FCLs in human keratinocytes. These epidermal cells are able to assemble different types of cell-matrix adhesions, including FAs and FCLs, and constantly need to adapt to changing conditions, such as coping with diverse mechanical forces. We report that binding of integrin αVβ5 to its ligand vitronectin in the presence of calcium is essential for the assembly of integrin αVβ5-containing FCLs. The clustering of integrin αVβ5 in FCLs is mediated by the clathrin adaptor proteins ARH and Numb that bind to the membrane proximal NPxY-motif in the integrin β5 cytoplasmic domain. Alternatively, Numb can also link integrin β5 and clathrin via an NPxY-motif independent mechanism through its interaction with EPS15/EPS15L1. Besides these mechanisms, we show that the localization of integrins in distinct cell-matrix adhesion complexes is controlled by actomyosin-driven cellular tension.

## Results

### Integrin αVβ5 localizes in FCLs in human keratinocytes

Here we analyze the distribution of integrin β5 in PA-JEB/β4 patient-derived immortalized keratinocytes in which the stable expression of integrin β4 was restored by cDNA transfection (Schaapveld et al., 1998). Using super-resolution microscopy (Fig. 1A) we show that integrin β5 is often found outside of FAs in circular structures. The morphology of these structures reminded us of the assembly of clathrin proteins, shown by EM as the irregular and triskelion structures that are located away from the cell periphery (Fig. 1B). By immunofluorescence and confocal microscopy imaging we confirmed that integrin β5 outside of FAs is concentrated on the ventral surface of the cells in clathrin-coated structures(Fig. 1C,D and 1SA). Morphometric analysis on integrin β5 clusters that are found outside FAs indicates that the shape and size of these clusters is very similar to that of FCLs (Fig. S1B), as previously described (Grove et al., 2014). These results were confirmed in HaCaT immortalized human keratinocytes, in which a similar subcellular distribution of integrin β5 can be observed (Fig. S1C). Integrin αV follows the distribution pattern of the β5 subunit (Fig. 1D), indicating the presence of heterodimerized αVβ5 in clathrin lattices. We do not find other integrin subunits associated with FCLs, as demonstrated for the RGD-binding integrin β1 (Fig. S1E). Occasionally, we can find integrin β5 in FCLs associated with actin (Fig. S1F), as has been previously described (Leyton-Puig et al., 2017).

**Fig. 1.**
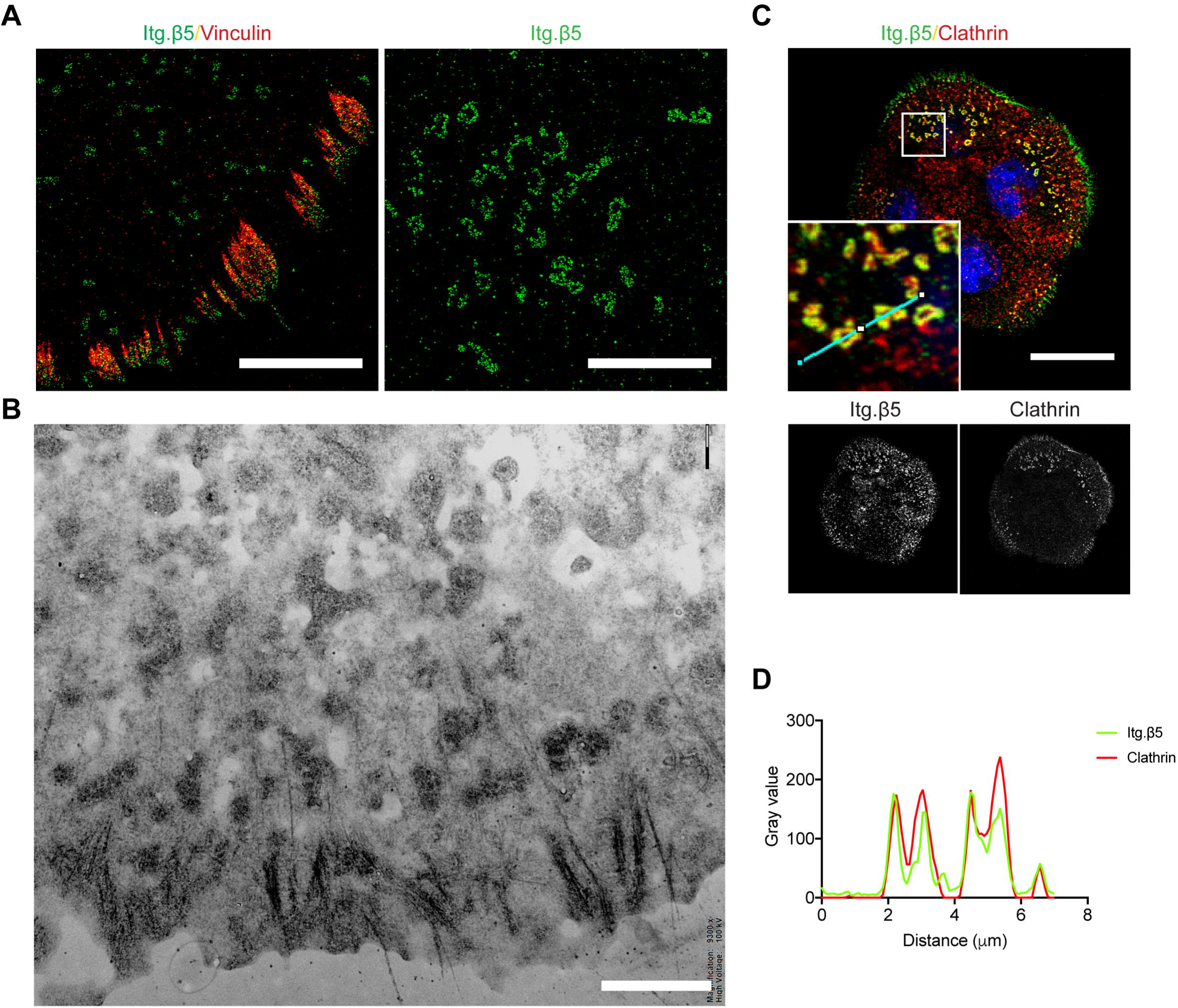
Integrin αVβ5 clusters are present in both focal adhesions and flat clathrin lattices in human keratinocytes. **A)** Representative super-resolution microscopy images showing integrin β5 (green) in and near focal adhesions, visualized by vinculin staining (red), at the cell periphery (left image) and more centrally located integrin β5 (right image). Scale bar, 5 μm. **B)** Electron microscopy image showing the area near the cell periphery containing both FAs and triskelion clathrin structures. Scale bar, 2 μm. **C)** Island of four keratinocytes showing integrin β5 (green in merge), clathrin (red in merge), and DAPI (blue). Scale bar, 20 μm. **D)** Intensity profiles of integrin β5 (green) and clathrin (red) along the cyan line in panel **C**.

### Clustering of integrin β5 requires vitronectin and calcium

The distribution pattern of integrin β5 as presented in Figure 1 could only be observed when the serum-free keratinocyte culture medium (KGM, containing epidermal growth factor and pituitary extract) had been replaced by DMEM supplemented with 10% fetal calf serum (FCS). To investigate whether vitronectin in the fetal calf serum-containing culture medium is critical for the formation of integrin β5-containing FCLs, we cultured keratinocytes on vitronectin-coated or uncoated coverslips in the presence of low or high calcium concentrations (KGM or DMEM culture medium, respectively). We observed that integrin β5-containing FCLs are only efficiently formed if both vitronectin and calcium are present (Fig. 2A), indicating that ligand binding, promoted by calcium (Asch and Podack, 1990), plays a role in integrin clustering. Similar results were obtained by culturing keratinocytes in calcium-depleted DMEM culture medium supplemented with low (0.09 mM) or high (1.8 mM) concentrations of calcium (Fig. S1G), which correspond with the calcium concentrations found in KGM or DMEM culture medium, respectively. Although the presence of vitronectin and calcium promotes the clustering of integrin β5 in FCLs, it does not alter its surface expression (Fig. 2B). Culturing keratinocytes on fibronectin-coated coverslips in the presence of a high calcium concentration did not lead to clustering of integrin β5 in FCLs (Fig. 2C). Integrin β5 clustering in the presence of vitronectin and calcium is needed for cell adhesion to vitronectin. Indeed, adhesion of keratinocytes to vitronectin in high calcium is significantly decreased upon treatment with cilengitide (Fig. 2D), an integrin αVβ3 and αVβ5 antagonist. Since like other normal keratinocytes, PA-JEB/β4 cells do not express the integrin β3 subunit ( data not shown, Adams and Watt, 1991; Kubo et al., 2001; Larjava et al., 1993), cilengitide could be considered as a specific inhibitor of integrin β5. Next, we studied whether integrin β5-containing FCLs behave as static or dynamic structures by preventing binding of integrin β5 to vitronectin in keratinocytes in which integrin β5 clusters had already been formed. Cilengitide acts as a competitive ligand-mimetic inhibitor but is not able to disrupt pre-existing integrin-ligand interactions (Mould et al., 2014), thus it can only prevent the formation of new integrin β5 clusters. We observed by immunofluorescence analysis that after 10 minutes of cilengitide treatment the integrin β5 clusters began to dissociate and after 30-90 minutes of treatment, only very few and small integrin β5 clusters remained visible at the basal membrane (Fig. 2E,F, Fig. S2A,C,E). However, cilengitide treatment did not decrease the surface expression of integrin β5 (Fig. S2D). Furthermore, cilengitide treatment did not result in a dramatic decrease of clustering of clathrin or of the clathrin adaptor protein Numb near the cell membrane (Fig. 2G, Fig. S2B,C,F), although the treatment resulted in slightly smaller clusters.

**Fig. 2.**
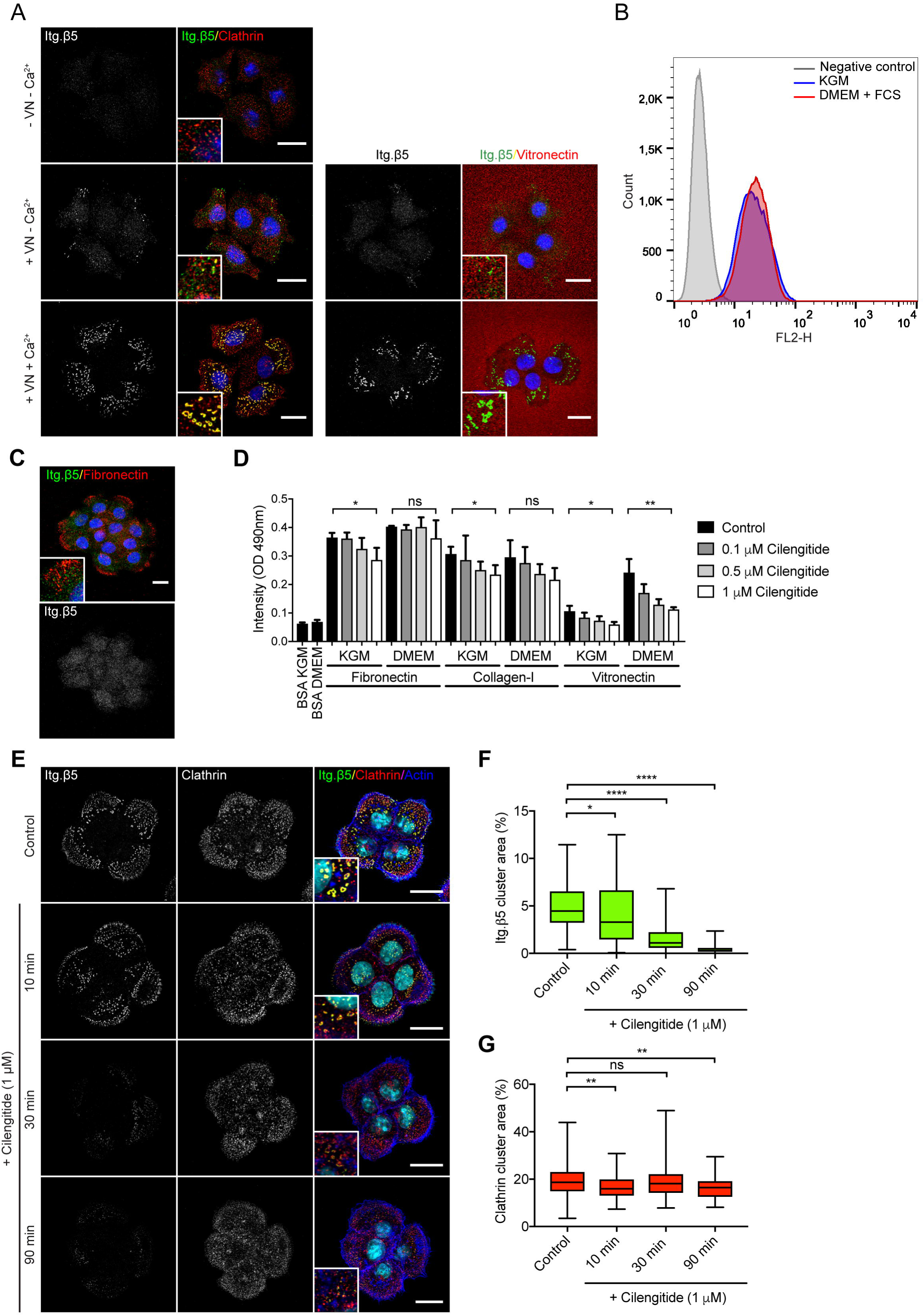
Binding to vitronectin in the presence of calcium is required for clustering of integrin β5 in flat clathrin lattices. **A)** Keratinocytes were grown on vitronectin-coated or uncoated coverslips in the presence or absence of high calcium levels. Images focus on the ventral cell surface. Left panel shows integrin β5 (green in merge), clathrin (red), and the cell nuclei (blue). Right panel shows vitronectin coating (red). **B)** FACS plot showing the expression of integrin β5 in keratinocytes grown in low calcium in KGM (blue) or in high calcium in DMEM supplemented with 10% FCS (red). Staining with the PE-conjugated secondary antibody only was used as a negative control (grey) (n=2). **C)** Keratinocytes seeded on fibronectin-coated coverslips. Integrin β5 (green in merge), fibronectin (red), and the cell nuclei (blue) are shown. **D)** Cells were treated with different concentrations of cilengitide (as indicated) in suspension, before a short-term (30 min) adhesion assay was performed on fibronectin-, collagen-, or vitronectin-coated substrates, in the presence (DMEM) or absence (KGM) of high calcium levels. Two-sided t-test was performed to calculate statistical significance between the control and samples treated with 1 μM cilengitide. *, P < 0.05; **, P < 0.01; ns, not significant. Columns show mean values with s.d. of three independent experiments. **E)** PA-JEB/β4 keratinocytes were grown in 10% FCS-supplemented DMEM culture medium overnight to induce integrin β5 clustering in FCLs and then treated with 1 μM cilengitide for the indicated times before fixation. Merged images show integrin β5 (green), clathrin (red), actin (blue) and the cell nuclei (cyan). **F,G)** The amount of integrin **(F)** or clathrin **(G)** clustering is defined as the total area of clusters on the cell membrane as a percentage of the total cell area. Data were obtained from three independent experiments. In total between 104 and 125 cells were analyzed per condition. Mann-Whitney U test was used to calculate statistical significance. *, P < 0.05; **, P < 0.01; ****, P < 0.0001; ns, not significant. Box plots range from the 25^th^ to 75^th^ percentile; central line indicates the median; whiskers show smallest to largest value. Scale bar, 20 μm.

### BioID reveals proximity interactors of integrin β5 clusters

To investigate what other proteins are associated with integrin β5-containing FCLs and might play a role in the recruitment of integrin β5 in clathrin lattices, we made use of the proximity biotinylation assay BioID (Roux et al., 2012). To this end, we deleted the integrin β5 subunit in PA-JEB/β4 keratinocytes using CRISPR/Cas9 and introduced an integrin β5-BirA* fusion protein, allowing the biotinylation of proteins in close proximity of the integrin β5 subunit. We performed BioID experiments in the presence or absence of calcium and vitronectin to detect proteins that are found associated with αVβ5 when it is concentrated in clathrin lattices. Identification of biotinylated proteins by mass spectrometry revealed multiple proteins involved in CME as proximity interactors of integrin αVβ5 (Fig. 3A, Table S1), including the adaptor protein complex 2 (AP2) subunits α1, β1, and μ1, epidermal growth factor receptor substrate15-like 1 (EPS15L1), intersectin-1 (ITSN1), and low density lipoprotein receptor adapter protein 1 (LDLRAP1 or ARH), and the E3 ligases Itch and NEDD4L. The presence of the proteins involved in endocytosis in the proximity of integrin β5 clusters was confirmed by immunofluorescence and calculation of Pearson’s correlation coefficient in PA-JEB/β4 and HaCaT keratinocytes (Fig. 3B-D, Fig. S3A). The cytoskeletal linker protein talin (TLN1) was also found by BioID, though immunofluorescence analysis shows that this protein is mainly present in FAs near the cell periphery and does not colocalize with integrin β5-containing FCLs (Fig. S3B).

**Fig. 3.**
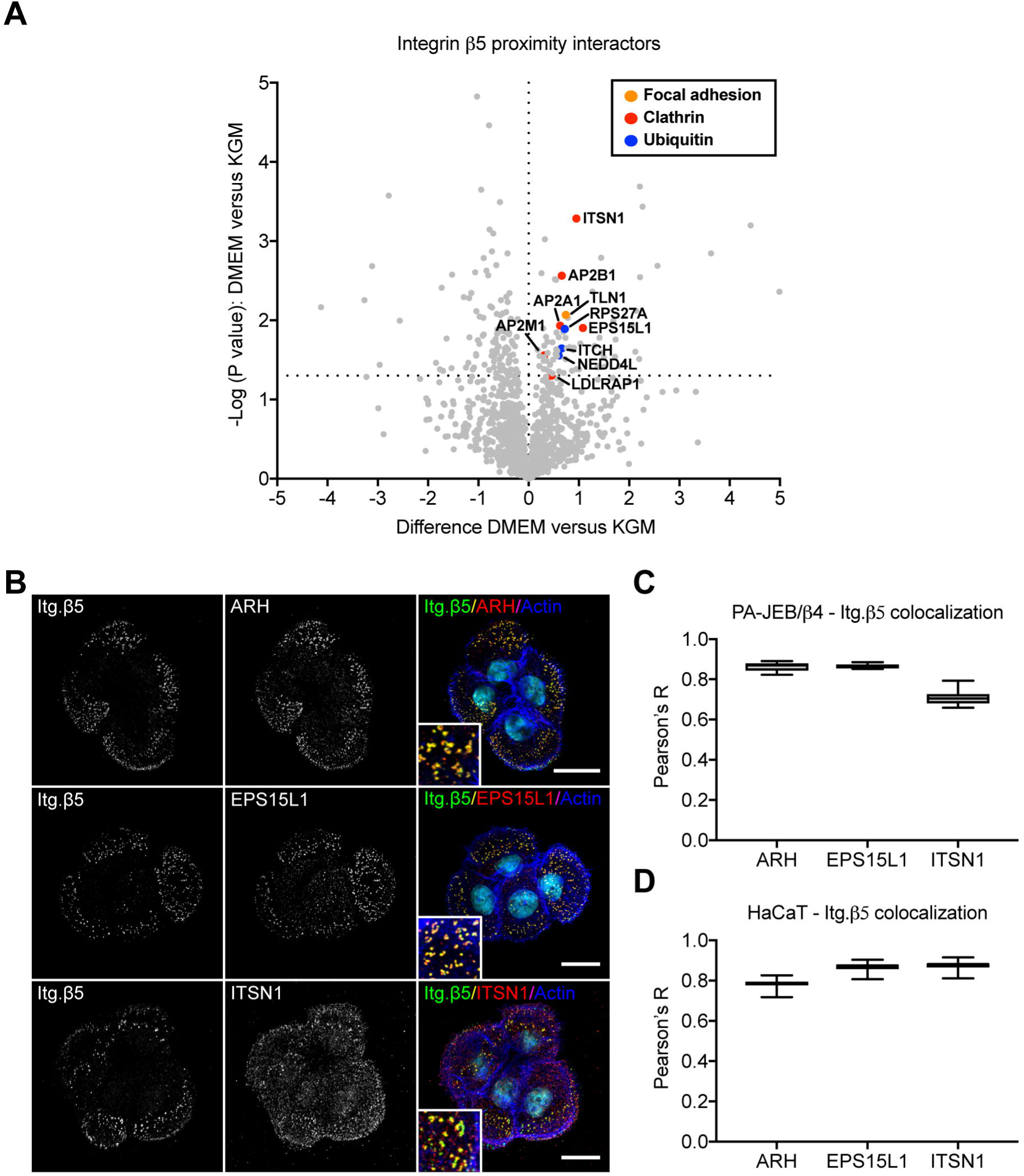
Clathrin adaptor proteins reside in close proximity of integrin β5 clusters. **A)** PA-JEB/β4 keratinocytes expressing integrin β5 fused to the promiscuous biotin ligase BirA* were used to perform proximity biotinylation assays with LC-MS/MS to determine the proximity interactors of integrin β5 subunits that are dispersed over the cell membrane (cell cultured in KGM) and of integrin β5 clusters (cells treated with 10% FCS-supplemented DMEM). Results of three independent experiments are shown in the volcano plot. The y-axis shows the negative log P values (dashed line at y=1.3 indicates a P value of 0.05) and the x-axis the difference in expression between the two conditions. Proteins are highlighted that are known to be associated with clathrin (red), FAs (orange), or play a role in protein ubiquitination (blue). Statistics: two-sided t-test. **B)** Representative confocal microscopy images (n=2) show the colocalization of integrin β5 (green in merge) and the clathrin adaptor proteins ARH, EPS15L1, and ITSN1 (red in merge). Nuclei are shown in cyan and actin in blue. Scale bar, 20 μm. **C)** Pearson’s correlation analysis of integrin β5 and ARH, EPS15L1, and ITSN1 in PA-JEB/β4 and in **D)** HaCaT keratinocytes. At least 24 cells obtained from 2 experiments were analyzed per condition. Box plots range from the 25^th^ to 75^th^ percentile; central line indicates the median; whiskers show smallest to largest value.

### ARH/Numb interact with the integrin β5 MP-NPxY motif

We wondered how the proximity interactors identified in the BioID screen interact with integrin β5 to regulate its localization in FCLs. The BioID assay does not distinguish between indirect and direct interactors of integrin β5. Since the latter ones are more likely to regulate the assembly of integrin β5-containing FCLs, we first analyzed which clathrin adaptor proteins can bind directly to the cytoplasmic domain of integrin β5. Integrin β5 has a short (<60 residues) cytoplasmic domain that contains a membrane-proximal (MP)-NPLY and membrane-distal (MD)-NKSY motif, two domains from which the NPxY and NxxY sequences are highly conserved among different integrin β subunits. These so called NPxY/NxxY motifs are canonical signals for clathrin-mediated endocytosis and serve as docking sites for PTB containing adaptor proteins (Calderwood et al., 2003; Traub and Bonifacino, 2013). The clathrin adaptor proteins ARH, protein disabled homolog 2 (Dab2), and Numb all contain a PTB domain (Mishra et al., 2002; Morris and Cooper, 2001; Santolini et al., 2000). PA-JEB/β4 keratinocytes express ARH and NUMB, but not Dab2 (Fig. S3C). ARH was identified in the BioID screen (Fig. 3A) and the presence of ARH in integrin β5-containing FCLs was confirmed by immunofluorescence analysis (Fig. 3B). We also found Numb associated with integrin β5-containing FCLs (Fig. S2C), although it had a lower significance in our BioID screen. To study the role of the NPxY motifs on clustering of integrin β5 in FCLs, we created mutants (Y>A) of the NPxY and NxxY motifs for integrin β5-BirA* in PA-JEB/β4 keratinocytes (Fig. 4A,B, Fig. S4A,B). In agreement with the notion that talin binds to the MP-NPxY motif and plays a key role in the formation of FAs, mutation of this motif but not that of the MD-NxxY motif, abrogated the localization of integrin β5 in FAs (Fig. 4A). The localization of integrin β5 in FCLs is not affected when either one or both NPxY and NxxY motifs have been mutated (Fig. 4B), indicating that the assembly of integrin β5-containing FCLs does not require the prior presence of the protein in FAs. Although integrin β5-containing FCLs were still able to form after mutating the NPxY and NxxY motifs, mutation of the MP-NPxY motif resulted in the formation of fewer and smaller, and more circular-shaped clusters (Fig. 4C-E). Next, we studied whether the mutations in the NPxY and NxxY motifs affected the proximity interaction between integrin β5 and ARH or Numb by using the BioID assay and analysis by western blot (Fig. 4F, Fig. S4C,D,E). Additionally, we studied colocalization of integrin β5 and ARH or Numb in FCLs by immunofluorescence and found that the MP-NPxY motif was required for the interaction of integrin β5 with ARH (Fig. 4G,H). A dispersed distribution of ARH was seen when binding of ARH to β5 was abrogated, yet ARH could still be observed in clathrin-coated structures (Fig. S4F). The MD-NxxY motif is not needed for the interaction between integrin β5 and ARH. Similar results were obtained for Numb (Fig. S5A,B), although the effect of the MP-NPxY mutation on the interaction between integrin β5 and Numb was less pronounced than that of integrin β5 and ARH. Finally, knock down of ARH reduced the number and size of integrin β5 clusters (Fig. 4I-K), indicating the role of this clathrin adaptor protein in regulating the localization of integrin β5 in FCLs.

**Fig. 4.**
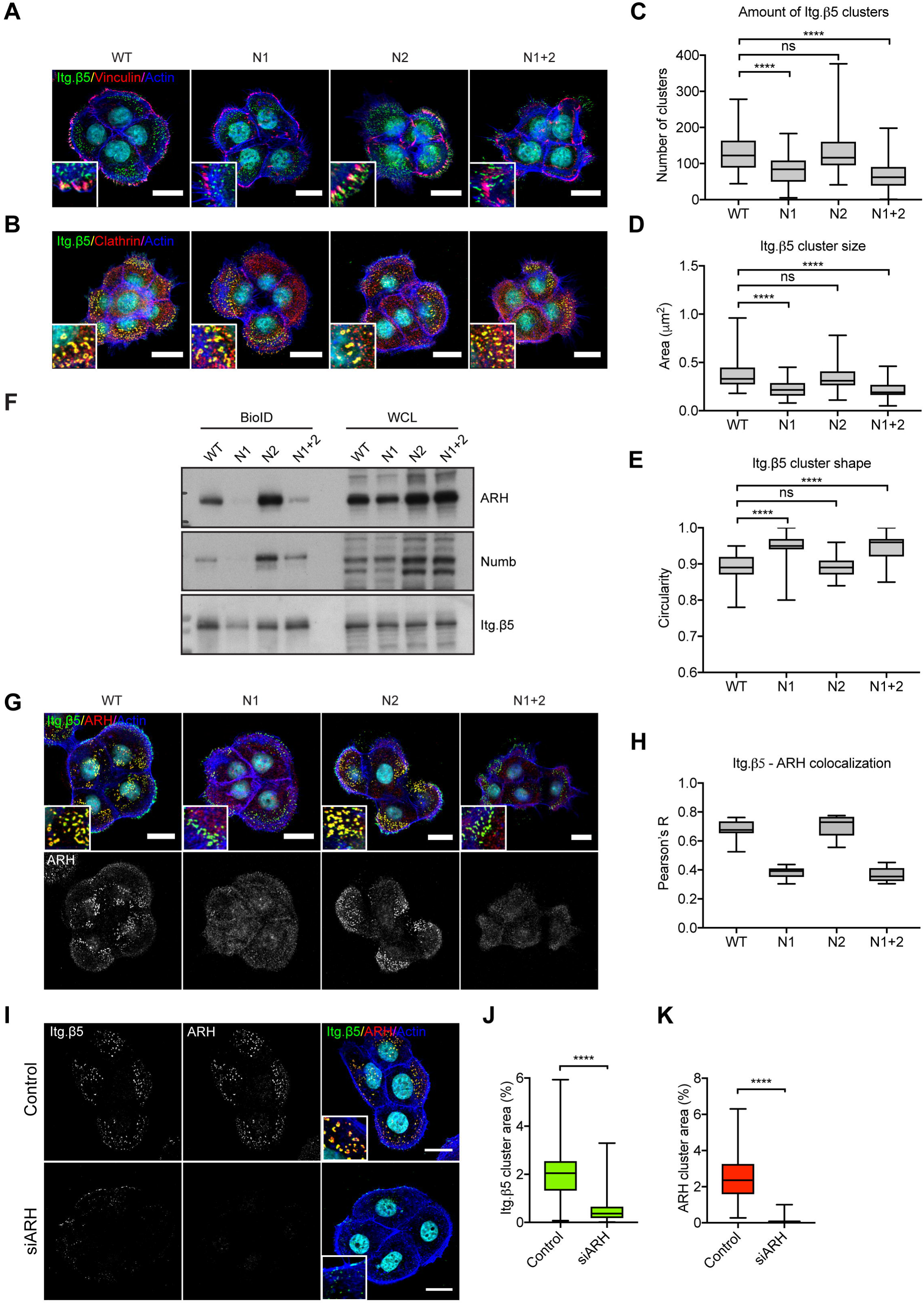
The membrane-proximal NPxY motif on the integrin β5 cytoplasmic domain interacts with the clathrin adaptor proteins ARH and Numb. **A)** Integrin β5 (green) does not cluster in FAs, visualized by vinculin staining (red) after mutation (Y>A) of the MP-NPxY motif (N1). See Fig. S4A for the sequence information. **B)** N1 and N2 Y>A mutations do not prevent clustering of integrin β5 (green) in clathrin structures (red). **C-E)** The number of integrin β5 clusters is reduced by N1 Y>A mutation and the average cluster size is reduced. The circularity of the smaller clusters as a result of the N1 mutant is increased. The y axis describes the shape ranging from 0 (irregular) to 1 (circle). At least 32 PA-JEB/β4 keratinocytes obtained from 2 independent experiments were analyzed per condition. Box plots range from the 25^th^ to 75^th^ percentile; central line indicates the median; whiskers show smallest to largest value. Scale bar, 20 μm. Mann-Whitney U test was performed to determine statistical significance. ****, P < 0.0001; ns, not significant. **F)** PA-JEB/β4 keratinocytes expressing integrin β5 (containing Y>A mutations in the MP-NPxY (N1) and/or MD-NxxY (N2) motif) fused to the promiscuous biotin ligase BirA* were used to perform proximity biotinylation assays. One representative western blot is shown out of three independent experiments. Quantifications of ARH/Numb signal intensities are shown in Fig. S4E. **G, H)** Proximity interaction between integrin β5 (green in merge) and ARH (red in merge) is reduced by the N1 Y>A mutation. ARH appears more diffuse over the cell membrane and the Pearson’s correlation coefficient is decreased. Scale bar, 20 μm. **I-K)** Knock down of ARH was accomplished by treating PA-JEB/β4 ARH siRNA for 24 h, prior to 24 h treatment with 10% FCS-supplemented DMEM culture medium to induce integrin β5 clustering. Merged images show integrin β5 (green), ARH (red), actin (blue) and the cell nuclei (cyan). Analysis of integrin β5 clustering shows a decrease of clustering after siRNA treatment. The amount of clustering is defined as the total area of clusters on the cell membrane as a percentage of the total cell area. Data were obtained from three independent experiments (approximately 100 cells in total). Mann-Whitney U test was performed to determine statistical significance. ****, P < 0.0001. Box plots range from the 25^th^ to 75^th^ percentile; central line indicates the median; whiskers show smallest to largest value. Scale bar, 20 μm.

### Numb/EPS15L1 bind β5 independently of the NPxY motif

Since mutating the NPxY motif does not completely abrogate integrin β5 clustering in FCLs, we investigated whether Numb could also mediate integrin β5 clustering in FCLs by a mechanism different from that of binding to the NPxY motif. Previous studies have shown that Numb binds EH-domain-containing proteins (Salcini et al., 1997), such as ITSN1 and EPS15L1, two scaffold proteins that are important for endocytosis by binding to clathrin and the AP-2 complex (Coda et al., 1998; Drake et al., 2000; Pechstein et al., 2010; Shih et al., 2002; Teckchandani et al., 2012; Wang et al., 2008; Yamabhai et al., 1998). Both proteins were identified as integrin β5 proximity interactors in our BioID screen (Fig. 3 and Table S1). EPS15L1 contains two ubiquitin-interacting motifs that are required for endocytosis of ubiquitinated cargo (Hofmann and Falquet, 2001), suggesting that this scaffold protein may form a link between clathrin and integrin β5 by binding to ubiquitin-modified β5. Indeed, integrin β5 is ubiquitinated, and its ubiquitination can be enhanced by inhibiting its lysosomal degradation or by preventing degradation of ubiquitinating enzymes by proteasomal inhibition (Fig. 5A). To test the hypothesis that an EPS15L1-Numb complex binds both ubiquitinated β5 and clathrin, we analyzed the effect of knock down of Numb and EPS15/EPS15L1 on integrin β5 clustering in FCLs. Depletion of Numb resulted in a significant decrease of β5 clustering in PA-JEB/β4 keratinocytes (Fig. 5B-D). Moreover, if Numb was depleted in keratinocytes expressing the β5 subunit carrying mutations in the NPxY and NxxY motifs, clustering of integrin β5 in FCLs could almost completely be prevented (Fig. 5B-D). Similar results were obtained for knock down of EPS15/EPS15L1 (Fig. 5E-G), providing evidence for the role of EPS15/EPS15L1 and Numb in localizing integrin β5 in clathrin lattices.

**Fig. 5.**
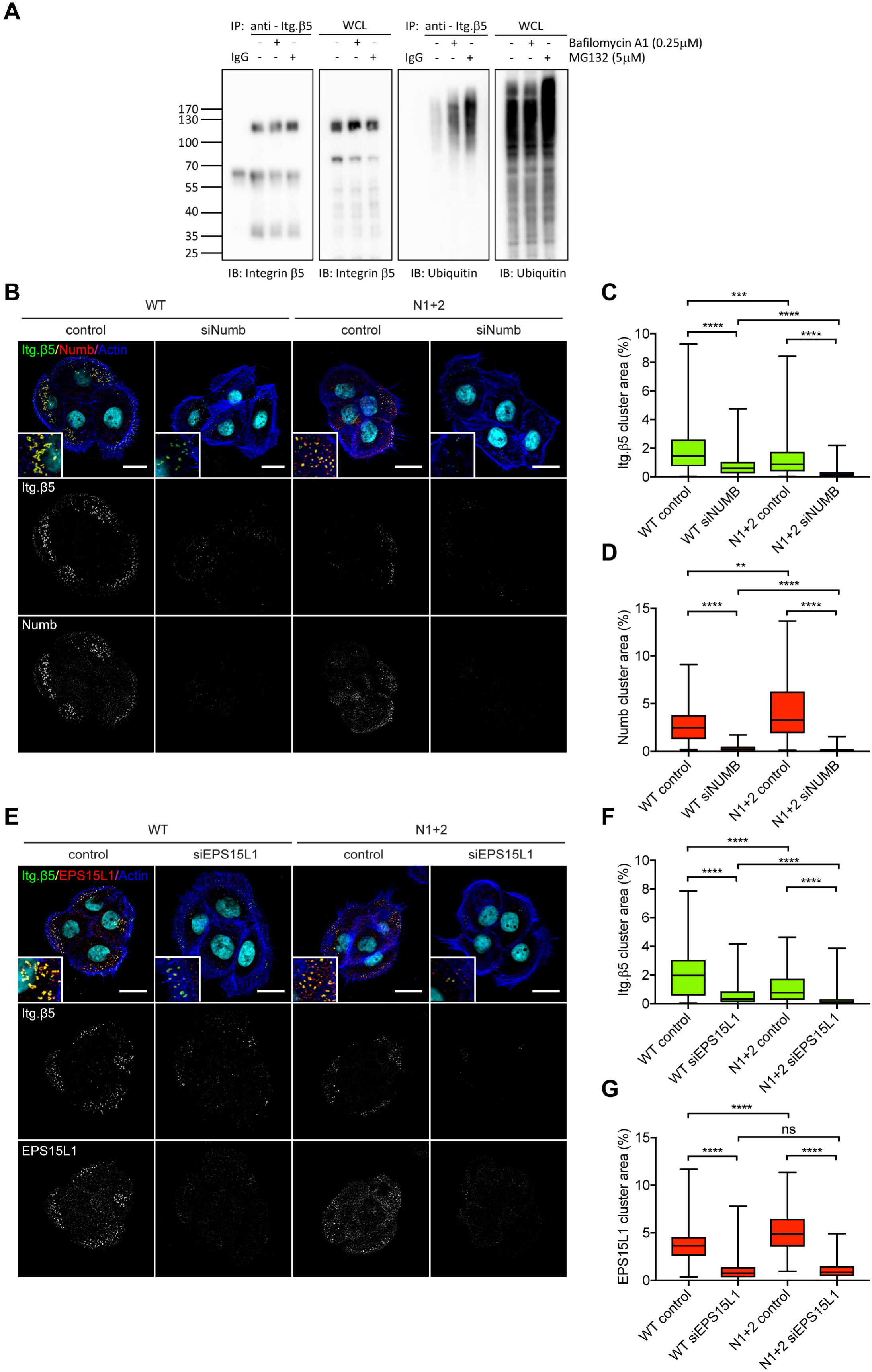
Knock down of Numb and EPS15/EPS15L1 prevents clustering of integrin β5 containing mutations in the NPxY motifs. **A)** Integrin β5 immunoprecipitation samples show (poly-) ubiquitination that is increased upon lysosomal or proteasomal inhibition, by treatment with 0.25 μM bafilomycin A1 and 5 μM MG132, respectively, for at least 3 h before cell lysis. Normal rabbit serum (IgG) is used as negative control for the immunoprecipitation. A representative western blot is shown (n=3). **B-G)** Knock down of Numb **(B)** or EPS15/EPS15L1 **(C)** was accomplished by treating PA-JEB/β4 keratinocytes (with Y>A mutations in the NPxY and NxxY motifs) with Numb or EPS15L1 and EPS15 siRNAs for 24 h, prior to 24 h treatment with 10% FCS-supplemented DMEM culture medium to induce integrin β5 clustering. Merged images show integrin β5 (green), Numb/EPS15L1 (red), actin (blue) and the cell nuclei (cyan). Analysis of integrin β5 clustering shows a decrease of clustering after siRNA treatment. The amount of clustering is defined as the total area of clusters on the cell membrane as a percentage of the total cell area. Data were obtained from three independent experiments (approximately 120 cells in total). Mann-Whitney U test was performed to determine statistical significance. **, P < 0.01; ***, P < 0.001; ****, P < 0.0001; ns, not significant. Box plots range from the 25^th^ to 75^th^ percentile; central line indicates the median; whiskers show smallest to largest value. Scale bar, 20 μm.

### Integrin β5 intracellular domain dictates clustering in FCLs

With the exception of integrin β5, RGD-binding integrins mainly cluster in FAs or FBs and associate with the actin cytoskeleton through an interaction with cytoskeletal linker proteins, such as talin. Integrin β5 and β3 are both RGD-binding integrins that share homologous regions (e.g. NPxY and NxxY motifs) and can both heterodimerize with the integrin αV subunit. Although PA-JEB/β4 keratinocytes do not endogenously express integrin β3, this integrin localizes in FAs and not in FCLs once exogenously introduced in these cells (Fig. S5C). Integrin β5 and β3 are both vitronectin receptors, however, integrin β3 binds many more ligands, including fibronectin, von Willebrand factor, thrombospondin, osteopontin, and fibrinogen (Humphries et al., 2006). We wondered whether differences in ligand binding are responsible for the localization of integrins in FCLs versus FAs and decided to create a system in which integrins possess equal ligand-binding properties and that enables us to study the contribution of the integrin β5 cytoplasmic domain on its localization in FCLs. To this end, we constructed different integrin chimeras that contain the extracellular and transmembrane domains of the β5 subunit and the intracellular domain of the β1 (β5^ex^/β1^in^) or β3 (β5^ex^/β3^in^) subunits and visualized their clustering by immunofluorescence (Fig. 6A,B). Like its full-length β3 counterpart, the integrin β5^ex^/β3^in^ chimera predominantly clustered in FAs (Fig. 6A,C) and less in FCLs (Fig. 6B,D). Similar results were obtained for the integrin β5^ex^/β1^in^ chimera, which is also found predominantly in FAs (Fig. 6). The distribution pattern of this chimera differs from that of its full-length counterpart (Fig. S1E), since the β5^ex^/β1^in^ chimera exclusively binds vitronectin, while the full-length β1 subunit can heterodimerize with 12 different α subunits and is thus able to reside in many different adhesion structures, including tetraspanin-enriched microdomains (Charrin et al., 2014; Hemler, 2005). A unique feature of the integrin β5 intracellular domain is the 8-amino acid stretch between the NPxY and NxxY motifs. However, deletion of this sequence did not prevent clustering of integrin β5 in FCLs (Fig. S5D,E). In summary, our results suggest that the intracellular domain of integrin β5 is important for its assembly in clathrin lattices.

**Fig. 6.**
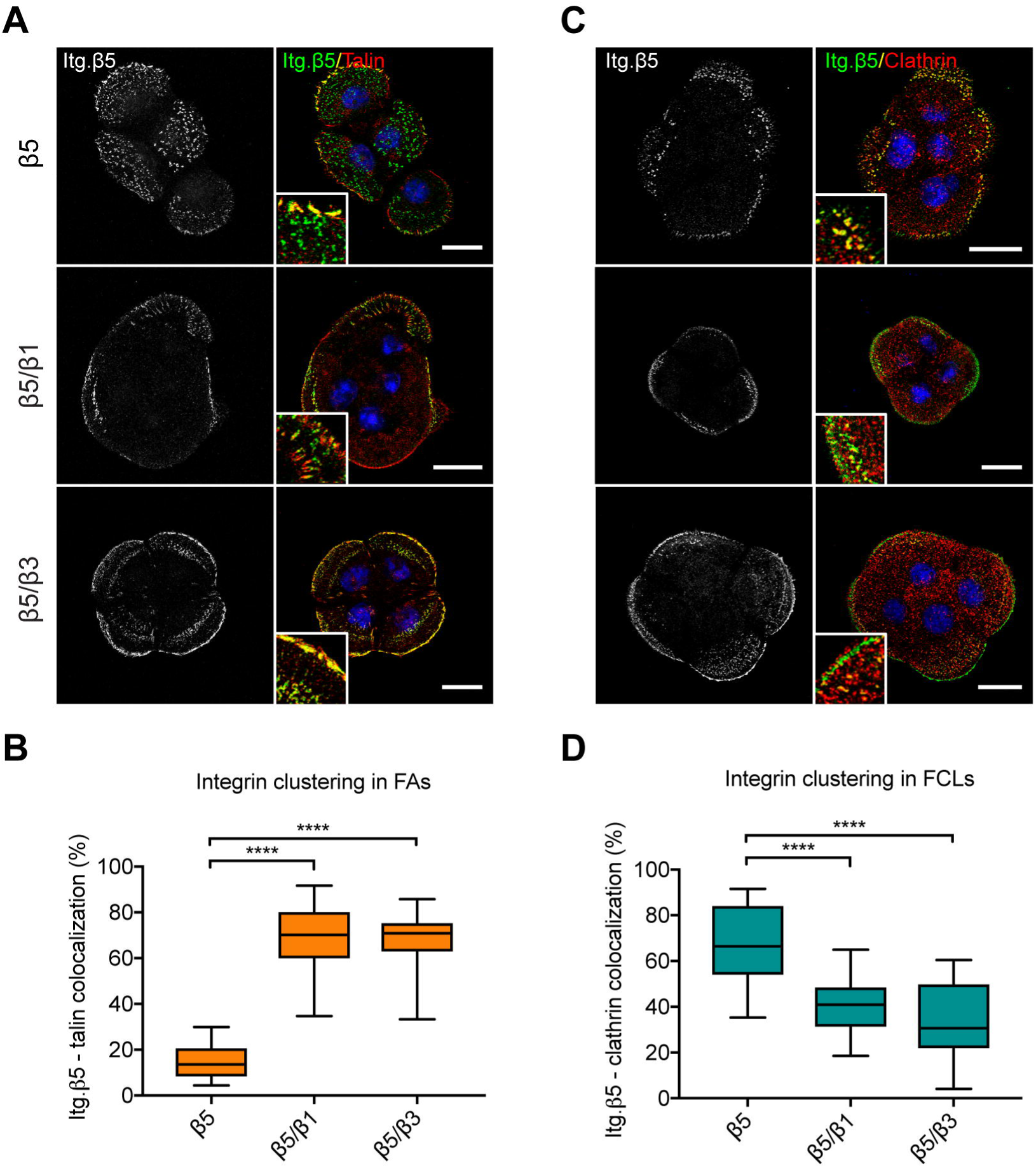
Integrin chimeras containing the intracellular domain of integrin β1 or β3 cluster predominantly in focal adhesions. **A)** Integrin β5, β5^ex^/β1^in^, and β5^ex^/β3^in^ (green in merge) colocalization with the FA marker talin (red). Nuclei are shown in blue. Occasionally, aspecific nuclear staining is detected in the integrin β5 channel. **B)** Integrin β5, β5^ex^/β1^in^, and β5^ex^/β3^in^ (green in merge) and clathrin structures (red). **C)** Integrin clustering in FAs defined by the area of integrin clusters overlapping with talin calculated as a percentage of the total integrin area per cell. **D)** Integrin clustering in FCLs defined by the area of integrin clusters overlapping with clathrin calculated as a percentage of the total integrin area per cell. Data were obtained from three independent experiments (n=30). Mann-Whitney U test was performed to determine statistical significance. ****, P < 0.0001. Box plots range from the 25^th^ to 75^th^ percentile; central line indicates the median; whiskers show smallest to largest value. Scale bar, 20 μm.

### Cellular tension regulates integrin localization

Although we did not find an association between exogenously expressed integrin β3 or the β5^ex^/β3^in^ chimera and clathrin, others have shown that the clathrin adaptor Dab2 binds to integrin β3 clusters when force generation is inhibited (Yu et al., 2015). In addition, the collagen-binding integrin α2β1 can be found in clathrin lattices when cells interact with collagen fibers in a soft 3D environment (Elkhatib et al., 2017). We hypothesized that the integrin subcellular distribution patterns might also be regulated by traction force generation. To this end, we treated keratinocytes with the myosin inhibitor blebbistatin in order to reduce cellular tension and observed that this treatment practically abolished clustering of integrin β5, β5^ex^/β1^in^, and β5^ex^/β3^in^ in FAs, as marked by vinculin staining (Fig. 7A,B) and favored their clustering in FCLs (Fig. 7C,D). A similar trend, though less dramatic, was observed by analyzing the colocalization of β5 and the different integrin β5 chimeras with the focal adhesion component talin and the clathrin adaptor protein Numb (Fig. S6).

**Fig. 7.**
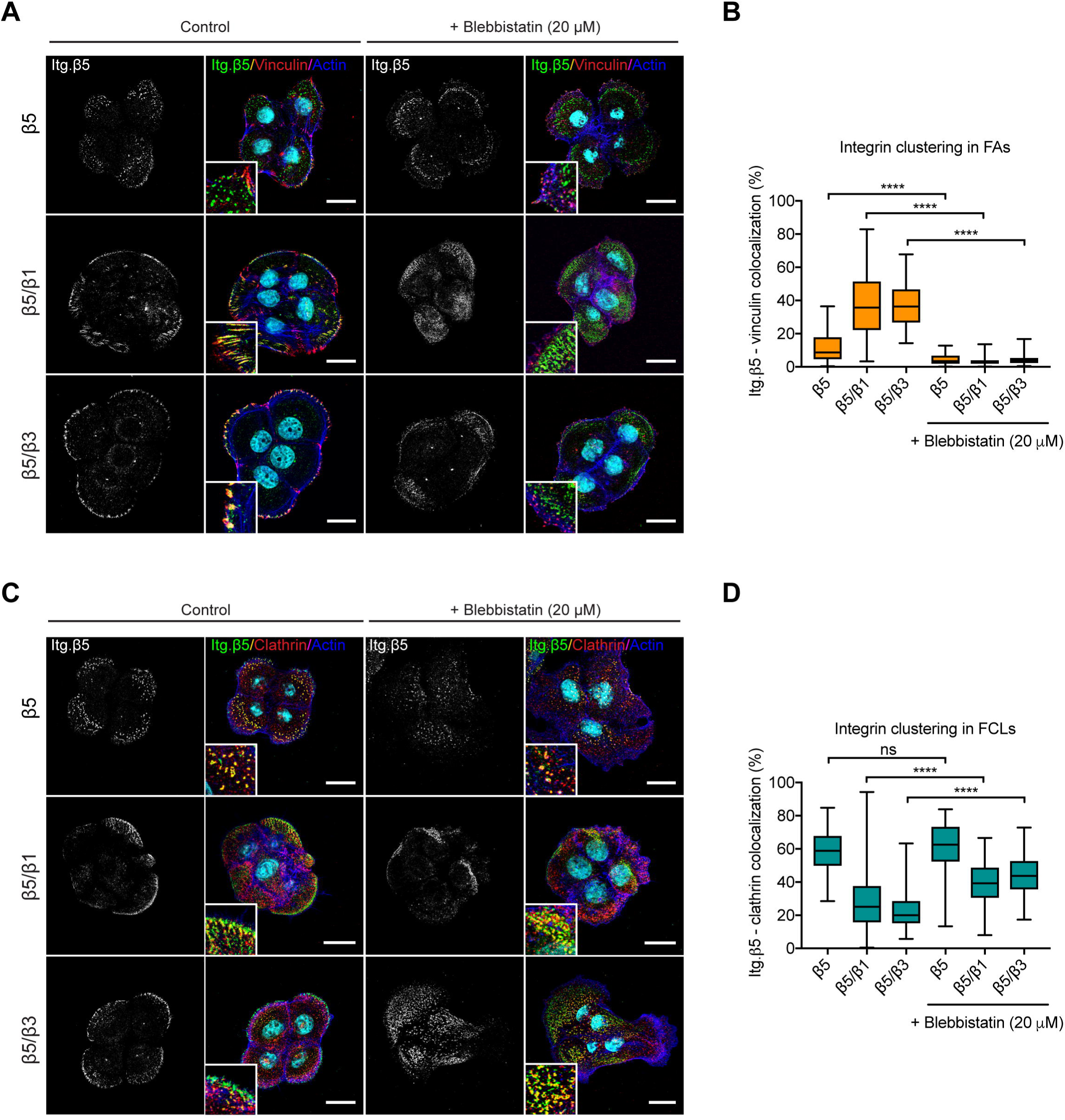
Integrin clustering is flat clathrin lattices versus focal adhesions is controlled by the amount of cellular tension. **A)** Integrin β5, β5^ex^/β1^in^, and β5^ex^/β3^in^ (green in merge) colocalization with the FA marker vinculin (red) is shown in response to treatment with the myosin inhibitor blebbistatin (20 μM) for 45 min prior to fixation. Actin is shown in blue and the nuclei in cyan. **B)** Integrin clustering in FAs is defined by the area of integrin clusters overlapping with vinculin calculated as a percentage of the total integrin area per cell. **C)** Colocalization of integrin β5, β5^ex^/β1^in^, and β5^ex^/β3^in^ (green in merge) with clathrin structures (red) with and without blebbistatin treatment. **D)** Integrin clustering in FCLs is defined by the area of integrin clusters overlapping with clathrin calculated as a percentage of the total integrin area per cell. Data were obtained from three biological replicates (60 cells total). Mann-Whitney U test was performed to determine statistical significance. *, P < 0.05; ****, P < 0.0001. Box plots range from the 25^th^ to 75^th^ percentile; central line indicates the median; whiskers show smallest to largest value. Scale bar, 20 μm.

Alternatively, we transfected PA-JEB/β4 keratinocytes with constitutively active RhoA to increase actomyosin contractility and cellular tension. We observed that in these keratinocytes the assembly of integrin β5-containing FCLs was decreased and that integrin β5 localized primarily in large FAs (Fig. 8). These findings were confirmed in HaCaT keratinocytes, in which integrin β5 colocalization with clathrin was decreased and colocalization with vinculin was increased after stimulation with LPA (Fig. S7). Clustering of integrin β5, either in clathrin lattices or focal adhesions, thus seems a dynamic and mechanosensitive process.

**Fig 8.**
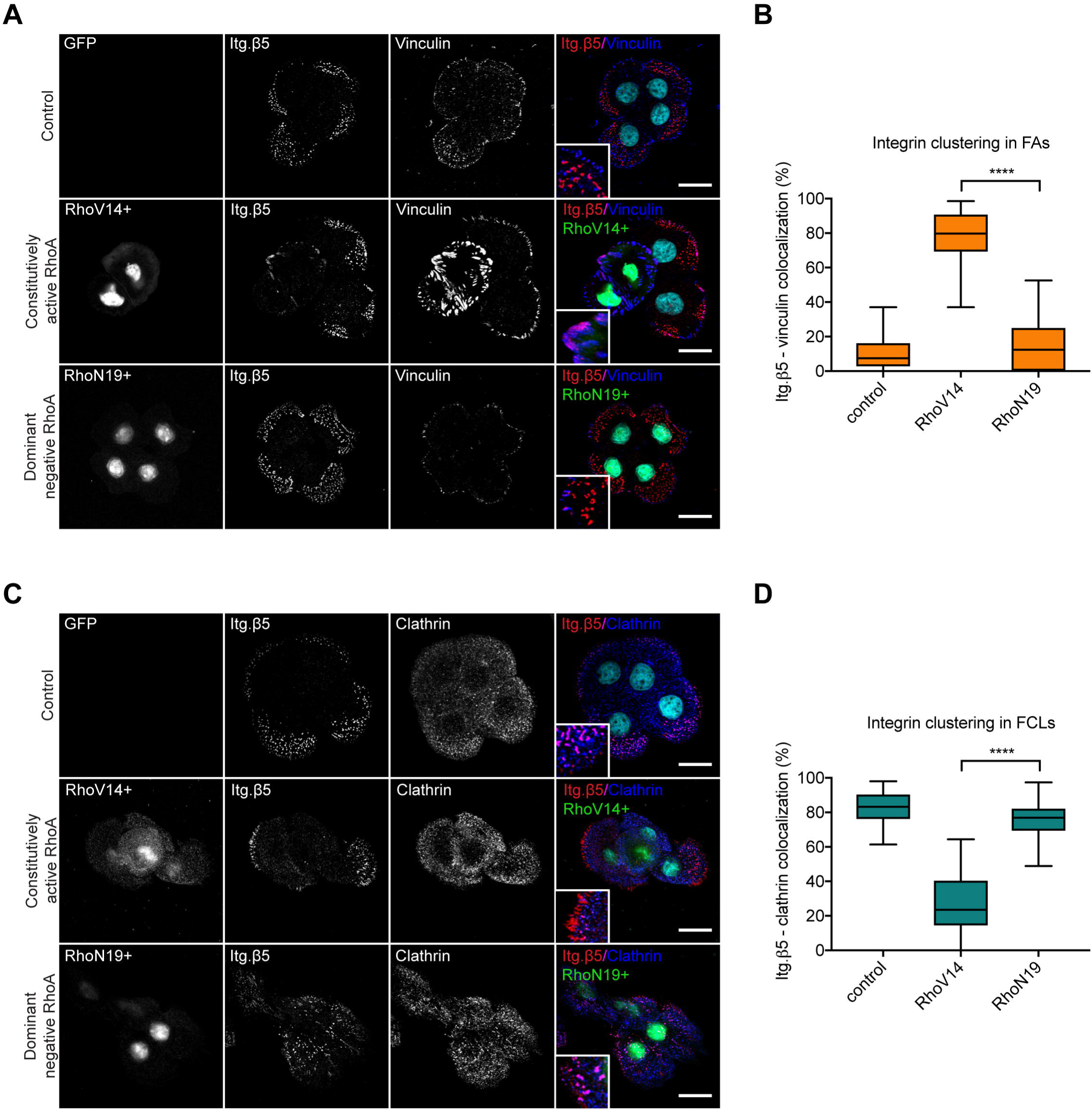
Increased cellular tension results in the clustering of integrin β5 in focal adhesions. **A+C)** PA-JEB/β4 keratinocytes were transiently transfected with constitutively active RhoA (V14) or dominant negative RhoA (N19) constructs. Transfected cells were selected based on the nuclear GFP signal. Integrin β5, vinculin/clathrin, and cell nuclei are shown in red, blue, and cyan, respectively, in the merged images. **B)** Integrin clustering in FAs defined by the area of integrin clusters overlapping with vinculin calculated as a percentage of the total integrin area per cell. **D)** Integrin clustering in FCLs defined by the area of integrin clusters overlapping with clathrin calculated as a percentage of the total integrin area per cell. Data were obtained from three independent experiments (60 cells total). Mann-Whitney U test was performed to determine statistical significance. ****, P < 0.0001. Box plots range from the 25^th^ to 75^th^ percentile; central line indicates the median; whiskers show smallest to largest value. Scale bar, 20 μm.

### Discussion

We investigated how the clustering of integrin αVβ5 in FCLs is regulated. Integrin αVβ5 is the only RGD-binding integrin that has been clearly shown to localize in these enigmatic clathrin structures, yet the reason of this distinct distribution pattern was so far unexplored. Here, we report that integrin αVβ5 clusters in FAs, but predominantly in FCLs, in human keratinocytes that are cultured in the presence of vitronectin and high levels of calcium (e.g. in serum-rich culture medium). The clustering of integrin αVβ5 in FCLs is a dynamic process. Blocking integrin-ligand interactions or increasing actomyosin-mediated cellular tension, results in reduced integrin αVβ5 clustering in FCLs.

The formation of FCLs in keratinocytes may be the result of “frustrated endocytosis”, triggered by integrin αVβ5 binding to vitronectin and the inability to internalize this serum protein because of its strong binding to the solid substratum. A recent study by Elkhatib *et al.* describes how the collagen binding integrin α2β1 associates with clathrin lattices to promote cell migration in soft 3D collagen gels (Elkhatib et al., 2017). Since fragments of collagen, generated by MMP cleavage, but not intact collagen fibers can be internalized by cells, the formation of integrin α2β1-containing clathrin lattices on collagen fibers can also be regarded as a form of frustrated endocytosis. This frustrated endocytosis of adhesion receptors might also lead to the “flat” organization of the clathrin lattices by preventing invagination of the membrane. FCLs may contribute to cell migration by providing additional sites of adhesion. In re epithelializing wounds, the binding of αVβ5 to vitronectin within the provisional matrix in the early wound bed may ensure that migrating keratinocytes remain associated with the substratum when exposed to mechanical forces. In line with the described findings that FCLs act as dynamic structures controlled by actin-based mechanisms (Leyton-Puig et al., 2017), our results demonstrate that integrin β5-containing FCLs rearrange in response to environmental changes, like the availability of ligand and/or the magnitude of force exerted on integrin αVβ5.

Noticeably, FCLs do not have to form as a result of frustrated endocytosis of integrin ligands per se. Large clathrin lattices can also be formed in the absence of adhesion molecules in skeletal muscles (Vassilopoulos et al., 2014) or at non-adherent cell surfaces (Grove et al., 2014). We propose that FCLs can be formed under different conditions and might be required for several biological processes. However, when integrins localize in these structures they act as alternative sites for cell-matrix adhesion.

Cell adhesion to vitronectin plays a major role in diverse biological processes. In normal keratinocytes, adhesion to vitronectin is solely mediated by integrin αVβ5 (Adams and Watt, 1991; Kubo et al., 2001; Larjava et al., 1993) and regulates keratinocyte migration and wound healing (Clark et al., 1996). Moreover, vitronectin synthesis and integrin β5 expression on tumor cells is correlated with disease progression of different tumor types, including melanoma (Desch et al., 2012; Vogetseder et al., 2013) and neural (Gladson et al., 1997; Uhm et al., 1999), breast (Bianchi-Smiraglia et al., 2013; Vogetseder et al., 2013) and non-small cell lung cancer (Bai et al., 2015; Vogetseder et al., 2013). Similar to keratinocytes, integrin β5 displays a punctate distribution over the ventral cell surface outside of FAs in human melanoma and lung carcinoma cells (Wayner et al., 1991). Integrin β5 interacts with different adaptor proteins in FCLs than in FAs (i.e. clathrin adaptor proteins versus cytoskeletal linker proteins). Therefore, localization of integrins in distinct adhesion structures might have important consequences for their downstream signaling and contribution to cellular processes that promote tumor progression. Furthermore, vitronectin regulates bacterial pathogenesis by serving as a cross-linker between bacteria and host cells and stimulating bacterial uptake through interaction with the host cell’s integrin receptors (Singh Molecular Microbiology 2010). The presence of integrin αVβ5 in FCLs might promote the uptake of pathogens and ECM components, due to the close proximity of this vitronectin receptor to the endocytic machinery and the notion that CME actively takes place at the edge of clathrin lattices (Lampe et al., 2016).

For the formation of integrin β5-containing FCLs, the intracellular domain of integrin β5 is crucial, as has been demonstrated previously (De Deyne et al., 1998; Pasqualini and Hemler, 1994) and shown in our studies with integrin chimeras. By making use of proximity-biotinylation assays (BioID), we were able to conduct unbiased screens to identify proximity interactors of integrin β5 in FCLs. We demonstrate that the clathrin adaptor proteins ARH and Numb interact with the MP-NPxY motif on the integrin β5 cytoplasmic domain. Moreover, an EPS15L1-Numb complex can interact with integrin β5 independently of the MP-NPxY motif and provides an additional mechanism for clathrin association, which is in line with other studies (Kang et al., 2013; Teckchandani et al., 2012). ARH does not contain the NPF motif that is needed for binding to EPS15L1, which could explain why the association between ARH and integrin β5 is lost after mutating the β5 MP-NPxY motif, while this mutation has a less dramatic effect on the association between integrin β5 and Numb. This observation is supported by the fact that knock down of Numb and EPS15/EPS15L1 has a larger effect on reducing the clustering of integrin β5 subunits carrying mutations in the NPxY and NxxY motifs than of wild type integrin β5, which can still interact with ARH. The interaction between EPS15L1-Numb and integrin β5 is likely to be mediated by binding of the ubiquitin-interacting motifs of EPS15L1 to ubiquitinated residues in the integrin β5 cytoplasmic domain. This interaction could possibly be enhanced by Numb, since this clathrin adaptor protein is able to further promote receptor ubiquitination by recruiting the E3 ligase Itch (Di Marcotullio et al., 2006; Di Marcotullio et al., 2011; McGill and McGlade, 2003). The clathrin adaptor Dab2 (not expressed in our cell lines) might have a similar role as Numb since it contains very similar domains (NPF) to bind EH-domain-containing proteins, like members of the epsin protein family.

Our data indicate that the integrin β5 localization pattern is regulated by the amount of cellular tension. When tension is increased, integrin β5 is localized predominantly in FAs. Alternatively, we show that integrin chimeras switch from clustering in FAs to FCLs when cellular tension is reduced. It is intriguing that under normal culture conditions the integrin β1, β3, and β5 subunits show a distinct subcellular distribution pattern, despite the presence of highly conserved domains. Since the β5^ex^/β1^in^ and β5^ex^/β3^in^ chimeras are able to cluster in FCLs under low cellular tension, it seems likely that the differences in integrin localization arise from the abilities of their β subunits to recruit specific adaptor proteins (Calderwood et al., 2003; Sun et al., 2016) and mediate traction forces. Indeed, it has been shown in pull down experiments that the β1, β3, and β5 subunit differ in their ability to bind kindlin-2 (Sun et al., 2016). We have confirmed these results and find that the β5 cytoplasmic domain is unable to bind kindlin-2 and kindlin-1 (unpublished). Hence, we surmise that on a stiff substratum the β5 integrin can mediate less traction forces than the β1 and β3 integrins, as it is unable to establish additional linkages to the actin cytoskeleton via kindlin and the ILK-PINCH-Parvin complex (Bledzka et al., 2016; Wickstrom et al., 2010). In apparent contrast to this view is that in cells subjected to high tension, the integrin β5 appeared to be primarily localized in large FAs rather than in FCLs. This pool of integrin β5, however, may not necessarily have to be involved in force generation but has become localized in FAs because of binding to the excessive amounts of talin available at these sites. In line with the fact that the β1 and β3 integrins by binding of their cytoplasmic domains to both talin and kindlin can mediate high traction forces on the substratum, we observed that the localization of the β5^ex^/β1^in^ and β5^ex^/β1^in^ chimeras in FAs is dramatically decreased when cellular tension is lowered. This is seen most clearly when FAs are marked by vinculin (Carisey et al., 2013; del Rio et al., 2009; Grashoff et al., 2010). When FAs were stained for talin, the decrease was less dramatic. This observation can be explained by the fact that talin, in contrast to vinculin, can reside in FAs under variable and low tension (Atherton et al., 2015; Kumar et al., 2016).

In agreement with our findings, integrin αVβ3 and α2β1 colocalize with clathrin in lattices in soft environments (Elkhatib et al., 2017; Yu et al., 2015). In general, it seems that integrin clustering in FCLs might give rise to additional sites of cell adhesion that are only formed when cells experience low tension. Yet, translocation to FAs occurs in response to increased actomyosin-based forces, which could be relevant in pathological conditions like cancer progression, in which vitronectin deposition and ECM stiffness are often increased.

In summary, we show how integrin αVβ5 clustering in FCLs is regulated by multiple mechanisms in the presence of vitronectin, high calcium, and low cellular tension. Our results provide novel insights into the regulation of integrin clustering in different adhesion structures. Additionally, these findings may shed light on the properties and function of the poorly characterized FCLs.

## Materials and Methods

### Antibodies

Primary antibodies used are listed in Table S2. Secondary antibodies were as follows: goat anti-rabbit Alexa Fluor 488, goat anti-mouse Alexa Fluor 488, goat anti-mouse Texas Red, goat anti-mouse Alexa Fluor 568, donkey anti-rabbit Alexa Fluor 594, goat anti-rabbit Alexa Fluor 647, and goat anti-mouse Alexa Fluor 647 (Invitrogen), PE-conjugated donkey anti-rabbit antibody (Biolegend #406421), stabilized goat anti-mouse HRP-conjugated and stabilized goat anti-rabbit HRP-conjugated (Pierce).

### Cell lines

Immortalized keratinocytes were isolated from a patient with Pyloric Atresia associated with Junctional Epidermolysis Bullosa (PA-JEB), as published elsewhere (Schaapveld et al., 1998). The derivation of this cell line was done for diagnostic purposes, thus the research conducted using these cells was exempt of the requirement for ethical approval. PA-JEB/β4 keratinocytes stably expressing integrin β4 were generated by retroviral transduction, as described previously (Sterk et al., 2000). Cells were maintained in serum-free keratinocyte medium (KGM; Invitrogen) supplemented with 50 μg ml^−1^ bovine pituitary gland extract, 5 ng ml^−1^ EGF, and antibiotics (100 units ml^−1^ streptomycin and 100 units ml^−1^ penicillin). HaCaT keratinocytes, obtained from the American Type Culture Collection were cultured in Dulbecco’s modified Eagle’s medium (DMEM) containing 10% heat-inactivated FCS and antibiotics. All cells were cultured at 37°C in a humidified, 5% CO_2_ atmosphere.

### Transient transfection

Human LDLRAP1 (M-013025-03-0020), EPS15 (M-004005-01-0020), EPS15L1 (M-004006-00-0020) and NUMB (M-015902-01-0020) siGENOME SMARTpool siRNAs were purchased from Dharmacon. The cDNAs encoding constitutively active (LZRS-IRES-GFP-RhoA(V14)) and dominant negative RhoA (LZRS-IRES-GFP-RhoA(N19)) were kindly provided by Jacques Neefjes. PA-JEB/β4 keratinocytes were transiently transfected with siRNAs or cDNAs using lipofectamine^®^ 2000 (Invitrogen). Lipofectamine (20 μl ml^−1^) and cDNA (6.5 μg ml^−1^) or siRNA (1 μM) solutions in Opti-MEM were mixed (1:1) and incubated for 20 min at room temperature. Cells were incubated with the transfection solution overnight.

### Generation of integrin β5-deficient keratinocytes

The target sgRNA against *ITGB5* (exon3; 5’-ACGGTCCATCACCTCTCGGT-3’) was cloned into pX330-U6-Chimeric_BB-CBh-hSpCas9 (a kind gift from Feng Zhang (Cong et al., 2013); Addgene plasmid #42230). PA-JEB/β4 keratinocytes were transfected with this vector in combination with a blasticidin cassette, as previously described (Blomen et al., 2015). Integrin β5-deficient cells were selected by supplementing the culture medium with 4 μg ml^−1^ blasticidin (Sigma) for four days following transfection.

### Stable cellular transduction

For the generation of the integrin β5-BirA* fusion proteins, pcDNA3-β5-BirA*(R118G) was obtained by inserting the coding sequence for β5, derived from pLenti-III-HA-ITGB5-mCherry (kindly provided by Dean Sheppard) together with the coding sequence of BirA* (R118G) into the *Eco*RI*/Xho*I sites of pcDNA3. The pcDNA3.1 MCS-BirA*(R118G)-HA plasmid was used as a template for the BirA* (a kind gift from Kyle Roux (Roux et al., 2012); Addgene plasmid #30647). Point or deletion mutants of β5 Y774 and/or Y794 and *del.8aa* were generated by site-directed mutagenesis with the PCR-based overlap extension method using Pwo DNA polymerase (Roche), and fragments containing the different mutations were exchanged with corresponding fragments in the β5 pcDNA3 vector. For the generation of the expression vectors encoding β5^ex^/β1^in^ and β5^ex^/β3^in^ chimeric integrin subunits, the codon encoding L744 in β5 was mutated from ctg to ctt, creating a *Hind*III site. Subsequently, this *Hind*III site was used for exchanging β1 or β3 cytoplasmic domain in the β5 pcDNA3 vector. Retroviral vectors containing mutant β5 cDNAs were generated by subcloning the mutant β5 cDNAs into the *Eco*RI and *Xho*I restriction sites of the LZRS-MS-IRES-Zeo vector. PA-JEB/β4 keratinocytes expressing different β5 mutants were generated by retroviral transduction. HEK293 cells were transiently transfected with 10 μg of cDNA, using the DEAE-dextran method.

### Super-resolution microscopy

For immunofluorescent analysis, keratinocytes grown on glass coverslips were fixed with 4% paraformaldehyde, permeabilized with 0.2% Triton-X-100, blocked in 5% BSA and incubated with primary and secondary antibodies at room temperature with extensive washing steps in between. Super-resolution microscopy was performed with a Leica SR GSD microscope (Leica Microsystems, Wetzlar, Germany) mounted on a Sumo Stage (#11888963) for drift free imaging. Collection of images was done with an EMCCD Andor iXon camera (Andor Technology, Belfast, UK) and a 160x oil immersion objective (NA 1.47). To image, the samples have been immersed in the multi-color super-resolution imaging buffer, OxEA (Nahidiazar et al., 2016). Laser characteristics were 405 nm/30 mW, 488 nm/300 mW and 647 nm/500 mW, with the 405 nm laser for back pumping. Ultra clean coverslips (cleaned and washed with base and acid overnight) were used for imaging. The number of recorded frames was variable between 10,000 to 50,000, with a frame rate of 100 Hz. The data sets were analyzed with the Thunder Storm analysis module (Ovesny et al., 2014), and images were reconstructed with a detection threshold of 70 photons, sub pixel localization of molecules and uncertainty correction, with a pixel size of 10 nm.

### Electron Microscopy (EM)

For EM, samples were fixed in Karnovsky’s fixative. Postfixation was done with 1% Osmiumtetroxide in 0.1 M cacodylatebuffer, after washing the cells were stained and blocked with Ultrastain 1 (Leica, Vienna, Austria), followed by ethanol dehydration series. Finally, the samples were embedded in a mixture of DDSA/NMA/Embed-812 (EMS, Hatfield, USA). This was all done in the tissue culture petri dish. Sectioning was performed parallel to the growing plane from the basal cell membrane upwards. Analysis was done with a Tecnai12G2 electron microscope (FEI, Eindhoven, The Netherlands).

### Flow cytometry

Cells were treated as indicated, trypsinized, and washed twice in PBS containing 2% FCS, followed by rabbit anti-human integrin β5 antibody (EM09902; 1:200 dilution) incubation for 1 h at 4°C. Then, cells were washed 3 times in PBS containing 2% FCS and incubated with PE-conjugated donkey anti-rabbit antibody (Biolegend #406421; 1:200 dilution) for 1 h at 4°C. After subsequent washing steps, integrin β5 expression was analyzed on a Becton Dickinson FACS Calibur analyser. For FACS sort, a PE-conjugated anti-human integrin β5 antibody (Biolegend #345203) was used and the desired cell populations were isolated using a Becton Dickinson FACSAria IIu or Beckman Coulter Moflo Astrios cell sorter.

### Adhesion assay

For adhesion assays, 96-well plates were coated with 10 μg ml^−1^ fibronectin from bovine plasma (Sigma #F1141), 5 μg ml^−1^ vitronectin (Sigma #SRP3186) or 3.2 μg ml^−1^ collagen 1 (Advanced Biomatrix #5005), overnight at 37°C. Before use, plates were washed once with PBS and blocked with 2% BSA (Sigma) in PBS for 1 h at 37°C. PA-JEB/β4 cells were trypsinized and resuspended in serum-free KGM or DMEM in the presence or absence of cilengitide (a kind gift from Coert Margadant). The cells were seeded at a density of 1×10^5^ cells per well and incubated for 30 min at 37°C. Nonadherent cells were washed away with PBS and the adherent cells were fixed with 4% paraformaldehyde for 10 min at room temperature, washed twice with H_2_O, stained with crystal violet for 10 min at room temperature and washed extensively with H_2_O. Cells were air-dried overnight and lysed in 2% SDS, after which absorbance was measured at 490 nm on a microplate reader using MPM6 software.

### Immunofluorescence

Unless mentioned otherwise, PA-JEB/β4 keratinocytes were seeded on glass coverslips and cultured in complete KGM medium for 24 h, and then treated with high calcium and vitronectin (DMEM + 10% FCS) for 24 h. HaCaT keratinocytes were seeded on glass coverslips and cultured in DMEM + 10% FCS. Cells were fixed with 2% paraformaldehyde for 10 min, permeabilized with 0.2% Triton-X-100 for 5 min, and blocked with PBS containing 2% BSA (Sigma) for at least 30 min. Next, cells were incubated with the primary antibodies for 1 h at room temperature. Cells were washed three times before incubation with the secondary antibodies for 1 h. Additionally, the nuclei were stained with DAPI and filamentous actin was visualized using Alexa Fluor 488 or 647-conjugated phalloidin (Invitrogen). After three washing steps with PBS, the coverslips were mounted onto glass slides in Mowiol. Images were obtained using a Leica TCS SP5 confocal microscope with a 63x (NA 1.4) oil objective.

### Image analysis and statistical analysis

Image analysis was performed using Fiji (ImageJ) (Schindelin et al., 2012; Schneider et al., 2012). For analysis of the colocalization between integrin β5 clusters and clathrin adaptor proteins, Pearson’s correlation coefficient was calculated (without threshold) using the JaCoP plug-in (Bolte and Cordelieres, 2006). Cluster size, amount, and circularity were calculated using the Analyze Particle function, after drawing a region of interest (ROI) at the cell periphery (based on actin staining). The total cluster area was divided by the total ROI area to define cluster density. To quantify integrin clustering in FAs (based on vinculin or talin staining) versus FCLs (clathrin staining), background was subtracted in both channels using a bilateral filter and the ROI was selected at the cell periphery. Colocalization of integrin clusters and FAs or FCLs was determined using the Image Calculator (command “multiply”) on both channels and calculating the area of overlapping clusters as a percentage of the total integrin cluster area per cell using the Analyze Particle function.

Mann-Whitney test (two-tailed P value) was performed using GraphPad Prism (version 7.0c). In figures, statistically significant values are shown as *, P < 0.05; **, P < 0.01; ***, P < 0.001; ****, p < 0.0001. Graphs were made in GraphPad Prism and show all data points represented as box-and-whisker plots, in which the box extends the 25^th^ to 75^th^ percentiles, the middle line indicates the median, and whiskers go down to the smallest value and up to the largest.

### BioID assay

PA-JEB/β4 expressing β5-BirA* cells grown on 145 mm plates were cultured in complete KGM or DMEM and treated with 50 μM biotin (Sigma #B4501) for 24 h. Cells were washed in cold PBS, lysed in RIPA buffer (20 mM Tris-HCl (pH 7.5), 100 mM NaCl, 4 mM EDTA (pH 7.5), 1% NP-40, 0.1% SDS, 0.5% sodium deoxycholate) supplemented with a protease inhibitor cocktail (Sigma), and cleared by centrifugation at 14.000 × *g* for 60 min at 4°C. Lysates were incubated with Streptavidin Sepharose High Performance beads (GE Healthcare) overnight at 4°C. Beads were washes three times with NP40 buffer (20 mM Tris-HCl (pH 7.5), 100 mM NaCl, 4 mM EDTA (pH 7.5), 1% NP-40) and twice with PBS and the isolated biotinylated proteins were analyzed by mass spectrometry or western blotting.

### Mass spectrometry

For mass spectrometry, samples were shortly separated on a 4-12% SDS-PAGE gel and stained with Coomassie Blue. The lane was excised from the gel after which proteins were reduced with dithiothreitol and alkylated with iodoacetamide. Proteins were digested with trypsin (mass spec grade, Promega) overnight at 37°C and peptides were extracted with acetonitrile. Digests were dried in a vacuum centrifuge and reconstituted in 10% formic acid for MS analysis. Peptide mixtures (33% of total digest) were loaded directly on the analytical column and analyzed by nanoLC-MS/MS on an Orbitrap Fusion Tribrid mass spectrometer equipped with a Proxeon nLC1000 system (Thermo Scientific). Solvent A was 0.1% formic acid/water and solvent B was 0.1% formic acid/80% acetonitrile. Peptides were eluted from the analytical column at a constant flow of 250 nl min^−1^ in a 140-min gradient, containing a 124-min linear increase from 6% to 30% solvent B, followed by a 16 min wash at 80% solvent B.

### Mass spectrometry data analysis

Raw data were analyzed by MaxQuant (version 1.5.8.3) (Cox et al., 2014) using standard settings for label-free quantitation (LFQ). MS/MS data were searched against the human Swissprot database (20,183 entries, release 2017_03) complemented with a list of common contaminants and concatenated with the reversed version of all sequences. Trypsin/P was chosen as cleavage specificity allowing two missed cleavages. Carbamidomethylation (C) was set as a fixed modification, while oxidation (M) was used as variable modification. LFQ intensities were Log2-transformed in Perseus (version 1.5.5.3) (Tyanova et al., 2016), after which proteins were filtered for at least two valid values (out of 3 total). Missing values were replaced by imputation based a normal distribution using a width of 0.3 and a downshift of 1.8. Differentially expressed proteins were determined using a t-test (threshold: P ≤ 0.05) and [x/y] > 0.3 | [x/y] < −0.3.

### Immunoprecipitation and western blotting

Subconfluent PA-JEB/β4 cells cultured in 100 mm cell plates were treated with DMEM supplemented with 10% FCS overnight and then additionally treated with 0.25 μM Bafilomycin A1 (InvivoGen #tlrl-baf1) or 5 μM MG132 (Sigma #M7449) for 3h at 37°C. Cells were washed in cold PBS and lysed in RIPA buffer (20 mM Tris-HCl (pH 7.5), 100 mM NaCl, 4 mM EDTA (pH 7.5), 1% NP-40, 0.1% SDS, 0.5% sodium deoxycholate) supplemented with 1.5 mM Na_3_VO_4_, 15 mM NaF (Cell Signaling), protease inhibitor cocktail (Sigma), and 5 mM *N*-methylmaleimide (Sigma). Lysates were incubated on ice for 10 min and cleared by centrifugation at 14.000 × *g* for 20 min at 4°C. Next, lysates were incubated with 1 μg ml^−1^ integrin β5 antibody (EM09902) or control normal rabbit serum for 2 h at 4°C. Subsequently, lysates were incubated for 2 h at 4°C with Protein G Sepharose 4 Fast Flow beads (GE Healthcare). Beads carrying the immune complexes were washed 4 times in RIPA buffer supplemented with inhibitors, eluted in sample buffer (50 mM Tris-HCl pH 6.8, 2% SDS, 10% glycerol, 12.5 mM EDTA, 0.02% bromophenol blue) containing a final concentration of 2% β-mercaptoethanol and denatured at 95°C for 10 min. Proteins were separated by electrophoresis using Bolt Novex 4–12% gradient Bis-Tris gels (Invitrogen), transferred to Immobilon-P transfer membranes (Millipore Corp) and blocked for at least 30 min in 2% BSA in TBST buffer (10 mM Tris (pH 7.5), 150 mM NaCl, and 0.3% Tween-20). Primary antibody (diluted 1:1000 in 2% BSA in TBST buffer) incubation took place overnight at 4°C. After washing twice with TBST and twice with TBS buffer, blots were incubated for 1 h hour at room temperature with horseradish peroxidase–conjugated goat anti-mouse IgG or goat anti-rabbit IgG (diluted 1:3.000 in 2% BSA in TBST buffer). After subsequent washing steps, the bound antibodies were detected by enhanced chemiluminescence using SuperSignal™ West Dura Extended Duration Substrate (ThermoFisher) or Clarity™ Western ECL Substrate (Bio-Rad) as described by the manufacturer. Signal intensities were quantified using ImageJ.

## Acknowledgements

We thank Bram van den Broek (Netherlands Cancer Institute) for his help with data analysis, and Marc Block, Simon Goodman, Marina Glukhova, Coert Margadant, and Jacques Neefjes for sharing reagents.

## Competing interests

The authors declare no competing interests.

## Funding

This work was supported by grants from the Netherlands Organization for Scientific Research (NWO; project number 824.14.010) and Dutch Cancer Society (project number 2013-5971) and by a grant from NWO as part of the National Roadmap Large-scale Research Facilities of the Netherlands, Proteins@Work (project number 184.032.201).

## Data availability

All relevant data are available from the authors on reasonable request.

## Supplementary Figure legends

**Supplementary Figure S1.** **A)** Montage of image slices making up a z-stack showing PA-JEB/β4 keratinocytes with integrin β5 (green in merge), clathrin (red in merge), actin (blue), and nuclei (cyan). Distance in z between the image slices is 1 μm. Scale bar, 20 μm. **B)** Morphometric analysis of integrin β5 clusters outside FAs (n=4). Circularity ranges from 0 (irregular) to 1 (circle). **C)** HaCaT keratinocytes showing integrin β5 (green in merge), clathrin (red in merge), actin (blue), DAPI (cyan). Scale bar, 20 μm. **D)** PA-JEB/β4 keratinocytes showing integrin β5 (green in merge), αV (red in merge), and DAPI (blue). **E)** PA-JEB/β4 keratinocytes showing integrin β5 (green in merge), β1 (red in merge), and DAPI (blue). **F)** Zoomed in regions of PA-JEB/β4 keratinocytes showing occasional overlap between integrin β5 clusters (green) and actin (grey). Scale bar, 5 μm. **G)** PA-JEB/β4 keratinocytes were grown in the presence of vitronectin and low or high calcium levels. Different calcium concentrations were obtained by first depleting calcium from FCS-supplemented DMEM culture medium using Chelex 100 resin, and then adding 0.09 mM (low) or 1.8 mM (high) CaCl_2_. Merged images show integrin β5 (green in merge), clathrin (red), actin (blue), and the cell nuclei (cyan). Scale bar, 20 μm.

**Supplementary Figure S2.** **A+E)** The number of integrin β5 clusters is reduced by cilengitide treatment and the average cluster size is reduced. The circularity of the smaller clusters as a result of cilengitide treatment is increased. The y axis describes the shape ranging from 0 (irregular) to 1 (circle). **B+F)** Number of clathrin **(B)** or Numb **(F)** clusters and their size and circularity. **C)** PA-JEB/β4 keratinocytes were grown in 10% FCS-supplemented DMEM culture medium overnight to induce integrin β5 clustering in FCLs and then treated with 1 μM cilengitide for the indicated times before fixation. Merged images show integrin β5 (green), Numb (red), actin (blue) and the cell nuclei (cyan). Scale bar, 20 μm. **D)** FACS plot showing the expression of integrin β5 in keratinocytes grown in DMEM supplemented with 10% FCS and treated with (red) or without (blue) 1 μM cilengitide for 90 min. Staining with the PE-conjugated secondary antibody only was used as a negative control (grey). Data were obtained from three independent experiments. In total between 104 and 125 cells were analyzed per condition. Mann-Whitney U test was performed to determine statistical significance. *, P < 0.05; ***, P < 0.001; ****, P < 0.0001; ns, not significant. Box plots range from the 25^th^ to 75^th^ percentile; central line indicates the median; whiskers show smallest to largest value.

**Supplementary Figure S3.** **A)** Representative confocal microscopy images show the colocalization of integrin β5 (green in merge) and the clathrin adaptor proteins ARH, EPS15L1, and ITSN1 (red in merge) in HaCaT keratinocytes. Nuclei are shown in cyan and actin in blue. **B)** PA-JEB/β4 keratinocytes showing integrin β5 (green in merge), talin (red in merge), actin (blue), and DAPI (cyan). Scale bar, 20 μm. **C)** Western blot showing the expression of the clathrin adaptor proteins Numb, ARH, and Dab2 in HaCaT and PA-JEB/β4 keratinocytes. HeLa cells were used as a positive control for Dab2 expression. GAPDH is used as loading control.

**Supplementary Figure S4.** **A)** Amino acid sequences of the cytoplasmic domain of wild type integrin β5 and the MP-NPxY and MD-NxxY mutants. The Y>A mutations are marked in red. **B)** Western blot showing the expression of integrin β5 wild type and MP-NPxY and MD-NxxY mutants fused to the BirA* biotin ligase (n=2). Fusion proteins were obtained by transfecting integrin β5-deficient PA-JEB/β4 keratinocytes (gRNA 2) as described in methods. **C,D)** Quantifications of signal intensities of ARH **(C)** and Numb **(D)** of three independent experiments. Bars show mean with s.d. **E)** Uncropped scans of the western blots shown in Fig. 4F. **F)** ARH (green in merge) appears more diffuse over the ventral cell membrane when the integrin β5 MP-NPxY motif is mutated. Colocalization with clathrin (red) is quantified using Pearson’s correlation coefficient (R). Actin is shown in blue and the nuclei in cyan. Scale bar, 20 μm.

**Supplementary Figure S5.** **A,B)** Proximity interaction between integrin β5 (green in merge) and Numb (red in merge) is reduced by the N1 Y>A mutation. Numb appears more diffuse over the cell membrane and the Pearson’s correlation coefficient is decreased. At least 32 PA-JEB/β4 keratinocytes obtained from 2 independent experiments were analyzed per condition. Box plots range from the 25^th^ to 75^th^ percentile; central line indicates the median; whiskers show smallest to largest value. Scale bar, 20 μm. **C)** Exogenously expressed integrin β3 (green in merge) colocalizes with talin (red) in integrin β5-deficient PA-JEB/β4 keratinocytes. Actin is shown in blue and the nuclei in cyan. Scale bar, 20 μm. **D)** Amino acid sequences showing the deletion of the 8-amino acid stretch in the cytoplasmic domain of integrin β5. **E)** Deletion of the 8-amino acid stretch does not prevent clustering of integrin β5 in FCLs. Merged image shows integrin β5 (green), clathrin (red), actin (blue) and the cell nuclei (cyan). Scale bar, 20μm.

**Supplementary Figure S6.** **A)** Integrin β5, β5^ex^/β1^in^, and β5^ex^/β3^in^ (green in merge) colocalization with the FA marker talin (red) is shown in response to treatment with the myosin inhibitor blebbistatin (20 μM) for 45 min prior to fixation. Actin is shown in blue and the nuclei in cyan. **B)** Integrin clustering in FAs is defined by the area of integrin clusters overlapping with talin calculated as a percentage of the total integrin area per cell. **C)** Colocalization of integrin β5, β5^ex^/β1^in^, and β5^ex^/β3^in^ (green in merge) with Numb structures (red) with and without blebbistatin treatment. **D)** Integrin clustering in FCLs is defined by the area of integrin clusters overlapping with Numb calculated as a percentage of the total integrin area per cell. Data were obtained from two independent experiments (30-50 cells total). Box plots range from the 25^th^ to 75^th^ percentile; central line indicates the median; whiskers show smallest to largest value. Scale bar, 20 μm.

**Supplementary Figure S7.** **A)** Integrin β5 (green in merge) colocalization with the FA marker vinculin (red in merge) is shown in response to treatment of HaCaT keratinocytes with 1 μM LPA for 60 min prior to fixation. Actin is shown in blue and the nuclei in cyan. **B)** Integrin clustering in FAs is defined by the area of integrin clusters overlapping with vinculin calculated as a percentage of the total integrin area per cell. **C)** Colocalization of integrin β5 (green in merge) with clathrin structures (red) with and without LPA stimulation. **D)** Integrin clustering in FCLs is defined by the area of integrin clusters overlapping with clathrin calculated as a percentage of the total integrin area per cell. Data were obtained from three biological replicates (60 cells total). Mann-Whitney U test was performed to determine statistical significance. ****, P < 0.0001. Box plots range from the 25^th^ to 75^th^ percentile; central line indicates the median; whiskers show smallest to largest value. Scale bar, 20 μm.

**Supplementary Table 2:** Primary antibody list.

| Antibody | Clone | Obtained from | Host | Application |
| --- | --- | --- | --- | --- |
| Integrin $\beta 5$ | EM09902 | Simon Goodman (Merck KGaA) | Rabbit | IF/FACS: 1:200<br>IP: $1\mu\text{g ml}^{-1}$ |
| Integrin $\beta 5$ | P1F6 | Biolegend (#920004) | Mouse | IF: 1:100 |
| Integrin $\beta 5$ | 5HK2 | Homemade | Rabbit | WB: 1:1000<br>SR: undiluted supernatant |
| Vinculin | VIIF9 | Marina Glukhova | Mouse | IF: 1:5 |
| Talin |  | Marc Block | Rabbit | IF: 1:100 |
| ARH/LDLRAP1 |  | AntibodyPlus (#A7093) | Rabbit | IF: 1:100<br>WB: 1:1000 |
| Numb | S.925.4 | Invitrogen<br>(#MA5-14897) | Rabbit | IF: 1:100<br>WB: 1:1000 |
| Clathrin, Heavy chain | X22 | Thermo Fisher<br>(#MA1-065) | Mouse | IF: 1:400 |
| ITSN1 |  | Atlas antibodies<br>(#HPA018007) | Rabbit | IF: 1:100 |
| EPS15L1 | EP1146Y | OriGene<br>(#TA301237) | Rabbit | IF: 1:100 |
| Ubiquitin | P4D1 | Covance<br>(#MMS-257P) | Mouse | WB: 1:1000 |
| Fibronectin | FN15 | Sigma<br>(SAB4200760) | Mouse | IF: 1:1000 |
| Vitronectin | 342603 | R&D Systems<br>(#MAB2349) | Mouse | IF: 1:1000 |
| Integrin β5-PE | AST-3T | Biolegend<br>(#345203) | Mouse | FACS |

